# All or Nothing: the False Promise of Anonymity

**DOI:** 10.1101/084921

**Authors:** Neil Walker

**Affiliations:** JDRF/Wellcome Trust Diabetes and Inflammation Laboratory, University of Oxford, Oxford, UK & Department of Clinical Informatics, University of Cambridge, Cambridge, UK

**Keywords:** anonymity, data sharing, de-identification, clinical trial

## Abstract

In early 2016, the International Committee of Medical Journal Editors (ICMJE) proposed that responsible sharing of de-identified individual-level data be required for clinical trials published in their affiliated journals. There would be a delay in implementing this policy to allow for the necessary informed consents to work their way through ethical review. Meanwhile, some researchers and policy makers have conflated the notions of de-identification and anonymity. The former is a process that seeks to mitigate disclosure risk though careful application of rules and statistical analysis, while the latter is an absolute state. The consequence of confusing the process and the state is profound. Extensions to the ICMJE proposal based on the presumed anonymity of data include: sharing unconsented data; sharing data without managing access, as Open Data; and proposals to sell data. This essay aims to show that anonymity (the state) cannot be guaranteed by de-identification (the process), and so these extensions to the ICMJE proposal should be rejected on governance grounds, if no other. This is not as negative a position as it might seem, as other disciplines have been aware of these limitations and concomitant responsibilities for many years. The essay concludes with an example from social science of managed access strategies that could be adopted by the medical field.

## Introduction

Funders have been seeking wider access to the research data they fund for many years (MRC 2000, OECD 2007). In the UK, in 2016 these aspirations have coalesced (RCUK 2016) and data sharing has become mandatory for some researchers (EPSRC 2014). However, in all cases funders’ policies make exceptions for confidential human subject data (University of Cambridge 2016).

In the context of clinical trials, the Alltrials campaign has been arguing for openness, with the slogan ‘All trials registered, all results reported’ (AllTrials 2013). But it falls short of demanding the sharing of individual patient data (IPD). OpenTrials (Goldacre and Gray 2016), the implementation arm of Alltrials, does not intend to include IPD, as it ‘often presents privacy risks that mean it cannot be simply posted online’ (OpenTrials 2016).

Recognising that sharing IPD should not be out of scope, since 2012, the Institute of Medicine’s (IOM) ‘Committee on Strategies for Responsible Sharing of Clinical Trial Data’ has been working on the theme, culminating in a January 2015 report ’sharing Clinical Trial Data: maximizing benefits, minimizing risks’ (IOM 2015).

In January 2016, the International Committee of Medical Journal Editors (ICMJE) responded with a widely-published proposal (Taichman et al. 2016) that requires authors to share with others the IPD underlying the results presented in the article.

They announced a year’s delay in implementation, to give time for the necessary ethical reviews and informed consents to be in place.

## From De-identification to Anonymity

While there were some protests at the ICMJE proposal, these were concerned with loss of scientific freedom implied (Lewandowsky and Bishop 2016) and included the controversial ‘research parasites’ editorial in NEJM (Longo and Drazen 2016). That this sort of data sharing in general, and de-identification in particular, was achievable was not questioned.

The ICMJE considers de-identification to be a way of mitigating the risk of disclosure in IPD from consenting trial participants. IOM, on the other hand, is happy to consider the sharing of unconsented data – where the data subjects were not asked about data sharing, rather than refused – if data can be de-identified ‘sufficiently’ (IOM 2015: 144). This implication was not lost on Dal-Ré (2016).

The long IOM Appendix B entitled ‘Concepts and Methods for De-identifying Clinical Trial Data’ cites and relies on the successful deposition of the International Stroke Trial (IST) database (Sandercock, Niewada and Członkowska 2011a, and see below]. This is an implementation of a widely published BioMed Central article (Hrynaszkiewicz et al. 2010), which discusses the preparation of raw clinical data for publication, by the redaction of direct and indirect identifiers. This is analogous to the US ’safe Harbor’ method of de-identifying protected health care information (HIPAA 2010).

While Hrynaskiewicz et al. (2010) recommend ‘consent for publication should be sought’, it is not considered essential, and the article chooses to focus instead on ‘confidentiality and anonymity’.

## The International Stroke Trial Database

The IST did not seek consent for data sharing from trial participants, and the de-identified data is available as Open Data from the University of Edinburgh data repository (Sandercock, Niewada and Członkowska 2011b).

The deposited data for IST has 19435 participants, and 112 variables. It is a very large trial, but some of the variable counts give cause for concern. For example, while there are 6257 participants from the UK, and 3437 from Italy, there are only 9 from Japan and 2 from France, a man and a woman.

Perhaps the concern is misplaced: the IST authors have removed all 28 direct and indirect identifiers referenced in Hrynaskiewicz et al. (2010) and there are millions of elderly men and women in France. However, while the checklist of direct and indirect identifiers does not include ‘Country’, the first indirect identifier has ‘Place of treatment or health professional responsible for care’. Therefore, the identity of the two French participants may only be protected by the fact that membership of the collaborative trial group was not published.

Subsequent guidance on anonymisation from the ICO makes the context-sensitive nature of indirect identifiers clearer (ICO 2012). In an anonymised dataset, data does not have to be aggregated into frequency records – a common misconception. Individual-level data records are permitted, and allowed to be unique, providing that the direct identifiers have been removed (name, address, DOB, NHS number etc., cf. HIPAA (2010)); and indirect identifiers are either removed or are put into classes that (taken in combination) reduce disclosure risk to an acceptably low level. An example may be to report age in age-bands, rather than as actual age; or location as the first part of the postcode rather than the whole.

Indirect identifiers depend on context. While gender is usually considered an unproblematic variable, in a dataset about breast cancers, the few male cases would stand out.

In order to define the context in which a variable is an indirect identifier, we need to answer the question ‘out of what set of people does a record have to be anonymous’? While tempting to answer ‘out of the whole population’, in a research setting the eligibility criteria and locations of subject recruitment are published, increasingly in Open Access journals. Therefore the upper bound for considering the anonymity of data records is that a record should be anonymous within the eligibility pool. And given that a subject’s participation in a study or trial might be known (e.g. through social media), a lower bound is that a record should be anonymous within the pool of subjects actually recruited.

As a footnote, the IST team advertised the availability of Open Data for a subsequent study, IST-3, in the Lancet (Sandercock et al. 2016a). However, on the repository website (Sandercock et al. 2016b) the data is embargoed until 2021, and is only available, if at all, to physical visitors ‘in order to comply with UK NHS Information Governance’.

## The Current State of Clinical Data Sharing

The IST authors are not alone.

Because there is a widespread belief in the value of open science and the possibility of anonymity, but little experience of managing access to controlled data, the current state of sharing clinical trial IPD can be characterised in Figure 1 (Strom et al. 2014; Bierer et al 2016).

**Figure 1:**
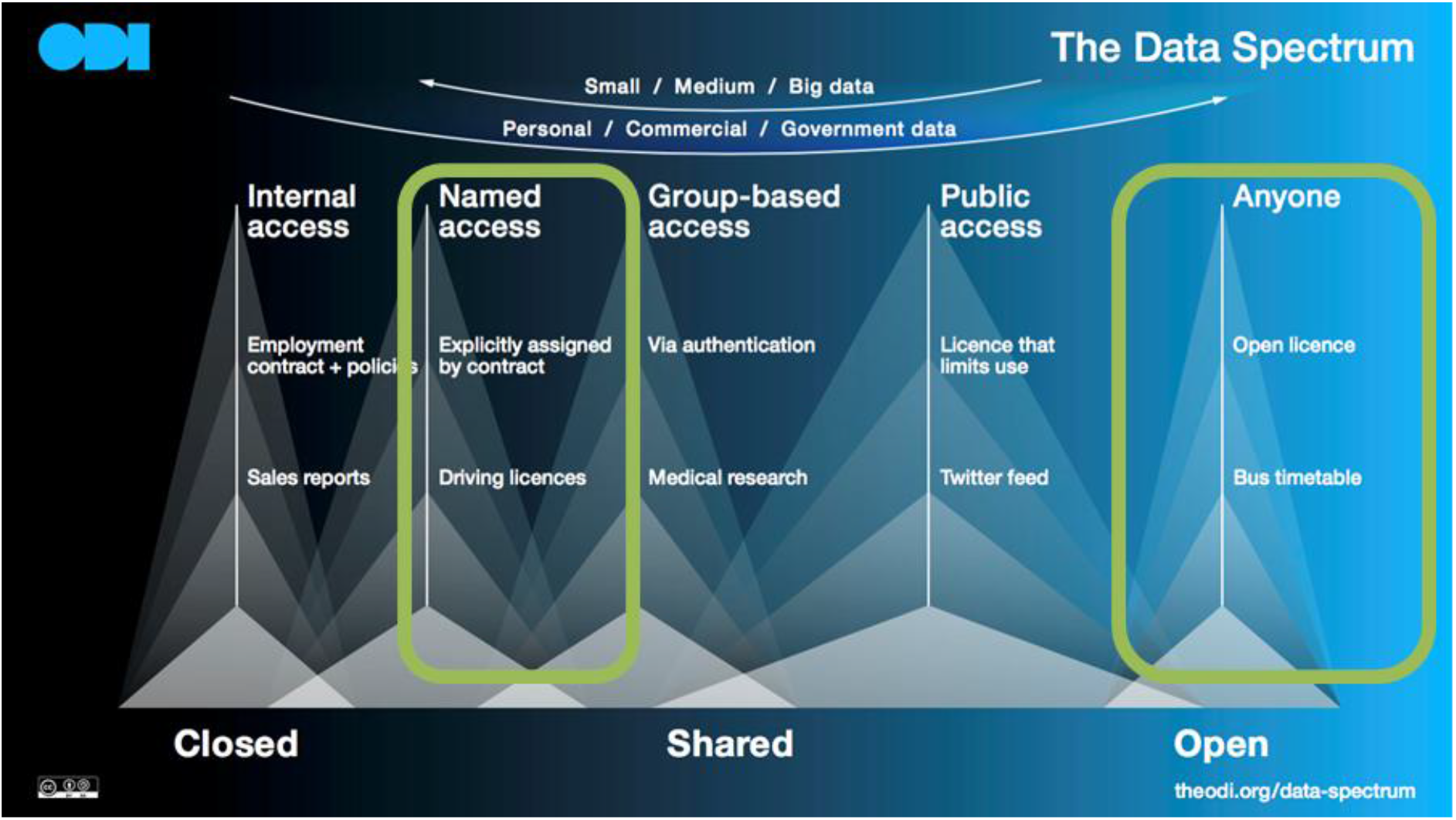
**The Open Data Institute’s (ODI) Data Spectrum**, showing the data sharing options most widely considered in clinical research. The ODI source image is CC-BY licensed.

This is problematic where the anonymity fails or falls under question. This happened when a potential forensic application of DNA matching in a pool of samples intruded into academic genetic research, and lead to the widespread removal of aggregate data from public websites (Zerhouni and Nabel 2008).

## Failures of Anonymisation

There have been many high-profile cases of failures in anonymisation, as investigated (and referenced) by Barth-Jones (2014). These failures can be split into four classes:

1. carelessness in redacting direct identifiers. An example is the Personal Genome Project distributing some sequence data in files named after the participants;
2. carelessness in redacting or grouping indirect identifiers. An example is the potential to identify men, by surname, from their Y-chromosome data, and of course, *care.data* (Carter, Lurie and Dixon-Wood 2015);
3. cases where the records are individually anonymous, but cease to be in combination. An example is the potential to identify mobile phone users by linking multiple locations;
4. cases where external knowledge can be linked to anonymous records to break that anonymity. While cases of linking media reports of prominent individuals to released datasets abound (e.g. NYC Taxi FOIL release, and health records in Iceland, UK and US), more subtle attacks exist – e.g. adding IMDb records to a Netflix release.

Failures in classes 1-3 are amenable to risk analysis. Failures in class 4 are less so, as we cannot know what other people know, nor the tools they will deploy. The ICO Anonymisation guide, puts it like this (ICO 2012: 16):

> You may be satisfied that the data your organisation intends to release does not, in itself, identify anyone. However, in some cases you may not know whether other data is available that means that re-identification by a third party is likely to take place.

The impact on participants should be noted. Whereas anonymisation advocates may be content with a small known residual risk (arrived at by testing for risks of class 1-3), the unknowability of risk of failure of class 4, and the widespread reporting of such failures, may persuade participants that research is not worthwhile.

## Alternatives to All or Nothing Approach

Once clinical trialists accept de-identification as a process of risk reduction, rather than a guarantor of a state of anonymity, they are free to adopt the risk assessment model that has been used so successfully by the social science community. In the UK, the UK Data Archive (UKDA 2016) holds such data, and its approach can be summarised in Figure 2.

**Figure 2:**
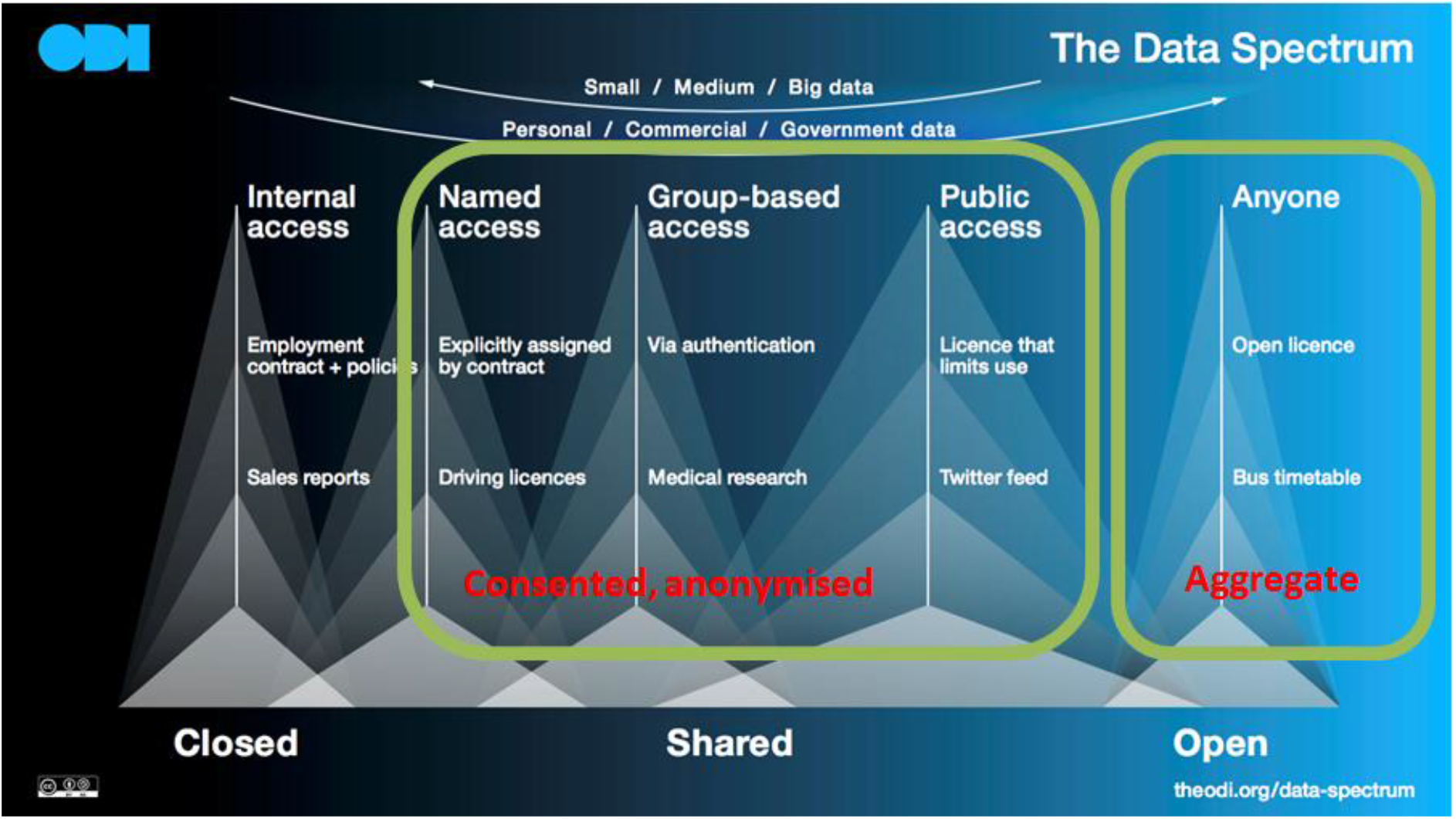
**The Open Data Institute’s (ODI) Data Spectrum**, showing the data sharing options used by the UK Data Archive. The ODI source image is CC-BY licensed.

Most (consented) individual-level data is distributed to registered users from registered host institutions using familiar website login mechanisms. But there is also provision to allow access to less heavily de-identified data, typically where some indirect identifiers remain, or where data is linked across sources, e.g. surveys linked to genetics. There is a range of data access mechanisms: by licence, where data is made available to a user who can supply the relevant credentials and has fulfilled the necessary training requirements; by application to data access committee, where a user presents a scientific justification for use of more identifiable data; and by use of a secure computing setting. These measures are combined: only a trained user with a scientifically approved proposal is allowed access to the secure computing resource. Usage of these measures is shown in Figure 3 for one of the major studies hosted at UKDA (Understanding Society 2016). Different versions of the same dataset, e.g. with geospatial identifiers included or redacted, may appear in several access categories, with the goal that the most heavily de-identified data is the most accessed. These access rates compare very favourably to reported rates from e.g. Strom et al. (2014). And as new identifiability threats are found, risks can be re-assessed, and data may be moved up and down the spectrum: the Homer et al (2008) problem (where aggregate data was shown to be potentially disclosive) would simply result in data moving from Anyone to Group-based Access, to use the ODI wording.

**Figure 3:**
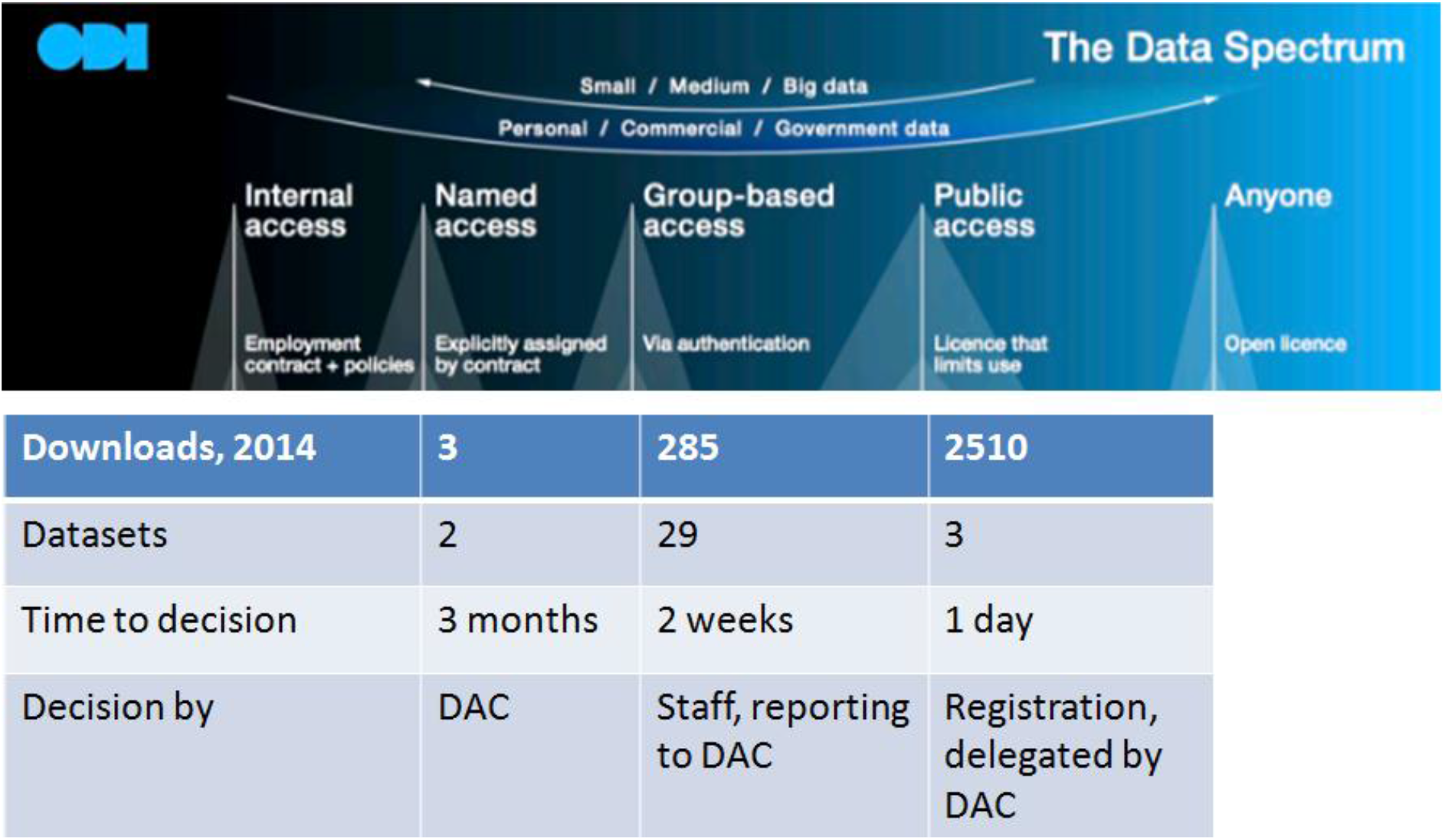
**The Open Data Institute’s (ODI) Data Spectrum**, with figures overlaid of successful access requests by the Understanding Society study group in 2014. The ODI source image is CC-BY licensed.

## Conclusion

There is a political agenda to de-identification – advocates claim it is possible, and that failures are the result of incompetence, a view that is heavily contested (Ohm 2009; Cavoukian and Castro 2014; Narayanan and Felten 2014).

There are incentives to make exaggerated claims of anonymity: to enhance openness, to reduce governance burden, but also to deny participants’ legitimate expectations and concerns.

Regardless of public expectation, the inability to guarantee anonymity should reinstate the ICMJE proposal in full: that de-identified IPD should be shared responsibly, though managed access, and in line with informed consent.

Finally, there are models beyond an All or Nothing approach, once the possibility of anonymity is dismissed and risk assessment is addressed seriously.

## Acknowledgements and Funding Statement

The views expressed are those of the author, and not those of the institutions to which he is and has been affiliated. I would like to thank members of the METADAC (Managing Ethico-social, Technical and Administrative issues in Data ACcess) data access committee (DAC) for many years of fruitful discussion; Jon Johnson for the formulation that we manage data access because we said we would; and Meena Kumari for the data from Understanding Society DAC. I would also like to thank my son, Owen Walker, for critical readings of both this essay, and the talk that preceded it.

The JDRF/Wellcome Trust Diabetes and Inflammation Laboratory is funded by the JDRF (9-2011-253), the Wellcome Trust [091157] and was funded by the National Institute for Health Research Cambridge Biomedical Centre, which funds the Department of Clinical Informatics. The Cambridge Institute for Medical Research (CIMR) is in receipt of a Wellcome Trust Strategic Award [100140]. The funders had no role in analysis, decision to publish, or preparation of the manuscript.

**All websites last accessed 31^st^ October 2016.**

## References

AllTrials 2013 What does all trials registered and reported mean? Webpage published September 2013. Available at http://www.alltrials.net/find-out-more/all-trials/

Barth-Jones, DC 2014 The antidote for “anecdata”: a little science can separate data privacy facts from folklore. Guest post at Harvard Info/Law, published 21^st^ November 2014. Available at https://blogs.harvard.edu/infolaw/2014/11/21/the-antidote-for-anecdata-a-little-science-can-separate-data-privacy-facts-from-folklore/

Bierer, BE, Li, R, Barnes, M and Sim, I 2016 A global, neutral platform for sharing trial data. N Engl J Med. 2016 Jun 23;374(25):2411–3. DOI: https://dx.doi.org/10.1056/NEJMp1605348

Carter, P, Laurie, GT and Dixon-Woods, M 2015 The social licence for research: why care.data ran into trouble. J Med Ethics. 41(5): 404–9. DOI: https://dx.doi.org/10.1136/medethics-2014-102374

Cavoukian, A and Castro, D 2014 Big data and innovation, setting the record straight: de-identification does work. Information and Privacy Commisioner, Ontario, Canada.

Dal-Ré, R 2016 The ICMJE trial data sharing requirement and participant’s consent. Eur J Clin Invest. DOI: https://dx.doi.org/10.1111/eci.12694

EPSRC 2014 Expectation – EPSRC website. Published from October 2014. Available at https://www.epsrc.ac.uk/about/standards/researchdata/expectations/

Goldacre, B and Gray J 2016 OpenTrials: towards a collaborative open database of all available information on all clinical trials. Trials. 17: 164. DOI: https://dx.doi.org/10.1186/s13063-016-1290-8

HIPAA 2010 Guidance Regarding Methods for De-identification of Protected Health Information in Accordance with the Health Insurance Portability and Accountability Act (HIPAA) Privacy Rule. Webpage updated from March 2010. Available at http://www.hhs.gov/hipaa/for-professionals/privacy/special-topics/de-identification/

Hrynaszkiewicz, I, Norton, ML, Vickers, AJ and Altman, DG 2010 Preparing raw clinical data for publication: guidance for journal editors, authors, and peer reviewers. BMJ, 340: c181. DOI: https://dx.doi.org/10.1136/bmj.c181 ICO 2012 Anonymisation: managing data protection risk code of practice. Published November 2012.

Available at: https://ico.org.uk/for-organisations/guide-to-data-protection/anonymisation/ The text excerpt is licensed from the Information Commissioner’s Office under the Open Government Licence (OGL) v3.0

IOM 2015 Sharing clinical trial data: maximizing benefits, minimizing risks. The National Academies Press. DOI: https://dx.doi.org/10.17226/18998

Lewandowsky, S and Bishop, D 2016 Research integrity: don’t let transparency damage science. Nature, 529(7587): 459–61. DOI: https://dx.doi.org/10.1038/529459a

Longo, DL and Drazen, JM 2016 Data sharing. N Engl J Med. 374(3): 276–7. DOI: https://dx.doi.org/10.1056/NEJMe1516564

MRC 2000 Personal Information in medical research. Published October 2000, revised January 2003. Available at https://www.mrc.ac.uk/documents/pdf/personal-information-in-medical-research/

Narayanan, A and Felten, EW 2014 No silver bullet: de-identification still doesn’t work. Published 9^th^ July 2014. Available at http://randomwalker.info/publications/no-silver-bullet-de-identification.pdf

OECD 2007 OECD principles and guidelines for access to research data from public funding. Published April 2007. Available at http://www.oecd.org/sti/sci-tech/38500813.pdf

Ohm, P 2009 Broken promises of privacy: responding to the surprising failure of anonymisation. In: UCLA L Rev 57: 1701

OpenTrials 2016 FAQ–OpenTrials. Available at http://opentrials.net/faq/

RCUK 2016 Concordat on Open Research Data launched. Published 28^th^ July 2016. Available at http://www.rcuk.ac.uk/media/news/160728/

Sandercock, PA, Niewada, M, Członkowska, A and International Stroke Trial Collaborative Group 2011 The International Stroke Trial database. Trials, 12: 101. DOI: https://dx.doi.org/10.1186/1745-6215-12-101

Sandercock, P, Niewada, M and Czlonkowska, A (2011) International Stroke Trial database (version 2), [dataset]. University of Edinburgh. Department of Clinical Neurosciences. DOI: http://dx.doi.org/10.7488/ds/104

Sandercock, P, Wardlaw, J, Lindley, R, Whiteley, W and Cohen, G 2016 IST-3 stroke trial data available. Lancet, 387(10031): 1904. DOI: https://dx.doi.org/10.1016/S1474-4422(16)30076-X

Sandercock, P, Wardlaw, J, Lindley, R, Cohen, G and Whiteley, W 2016 The third International Stroke Trial (IST-3), 2000-2015 [dataset]. University of Edinburgh & Edinburgh Clinical Trials Unit. DOI: https://dx.doi.org/10.7488/ds/1350

Strom, BL, Buyse, M, Hughes, J and Knoppers, BM 2014 data sharing, year 1 – access to data from industry-sponsored clinical trials. N Engl J Med. 371(22): 2052–4. DOI: https://dx.doi.org/10.1056/NEJMp1411794

Taichman, D.B, Backus, J, Baethge, C, Bauchner, H, de Leeuw, PW, Drazen, JM, Fletcher, J, Frizelle, FA, Groves, T, Haileamlak, A, James, A, Laine, C, Peiperl, L, Pinborg, A, Sahni, P and Wu, S 2016 PLoS Med. 13(1): e1001950. DOI: https://dx.doi.org/10.1371/journal.pmed.1001950

UKDA 2016 UK data archive – home. Available at http://www.data-archive.ac.uk/

Understanding Society 2016 Understanding Society data access strategy v22. Published February 2016. Available at https://www.understandingsociety.ac.uk/documentation/getting-started

University of Cambridge 2016 Funders’ Policies | Research Data Management. Updated since 2014. Available at http://www.data.cam.ac.uk/funders

Zerhouni, EA and Nabel, EG 2008 Protecting aggregate genomic data. Science. 322(5898): 44. DOI:https://dx.doi.org/10.1126/science.1165490

